## Supplementary figures and images for "Developmental stage-specific spontaneous activity contributes to callosal axon projections"

### Figure 1-figure supplement 1

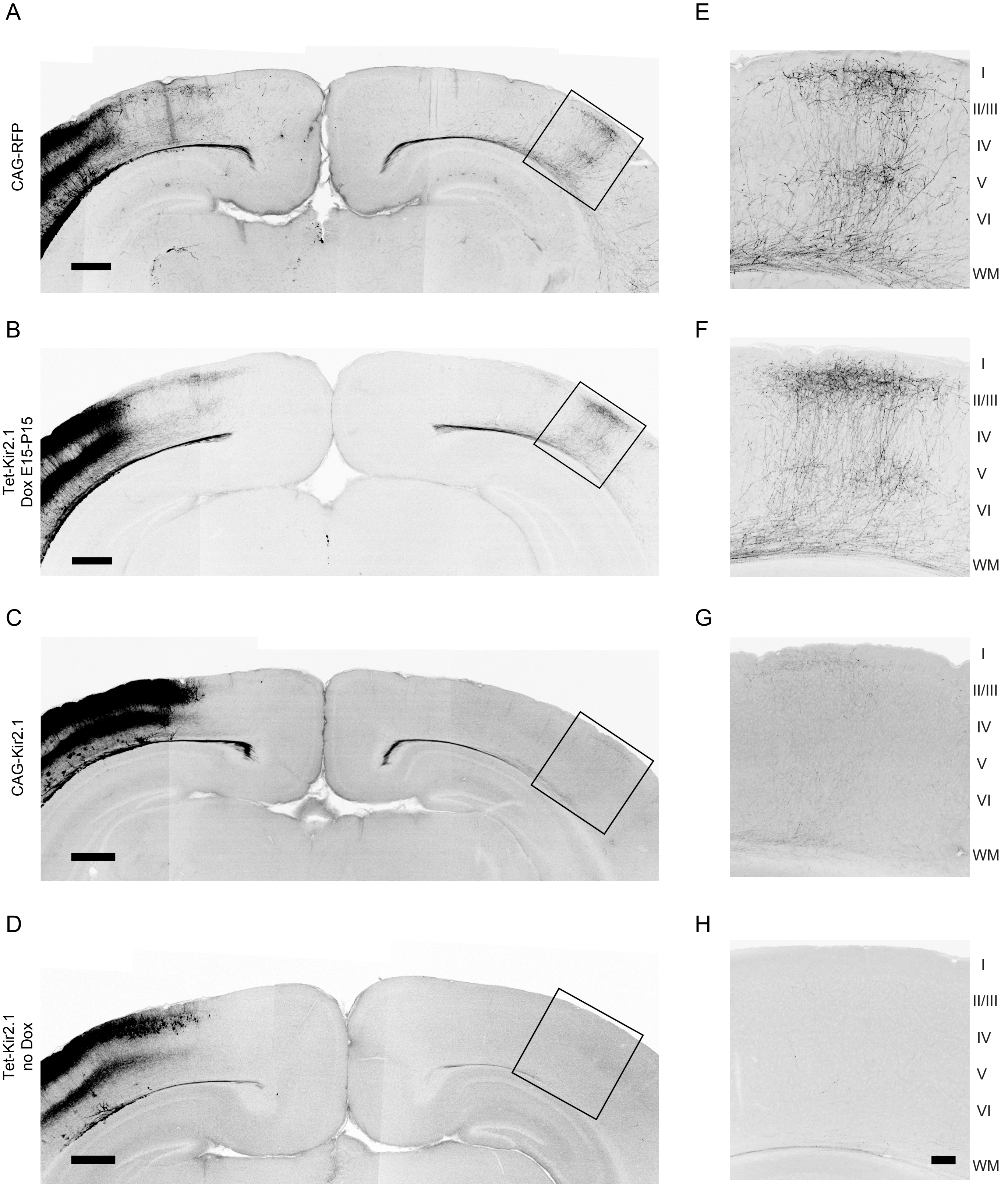

### Figure 1-figure supplement 2

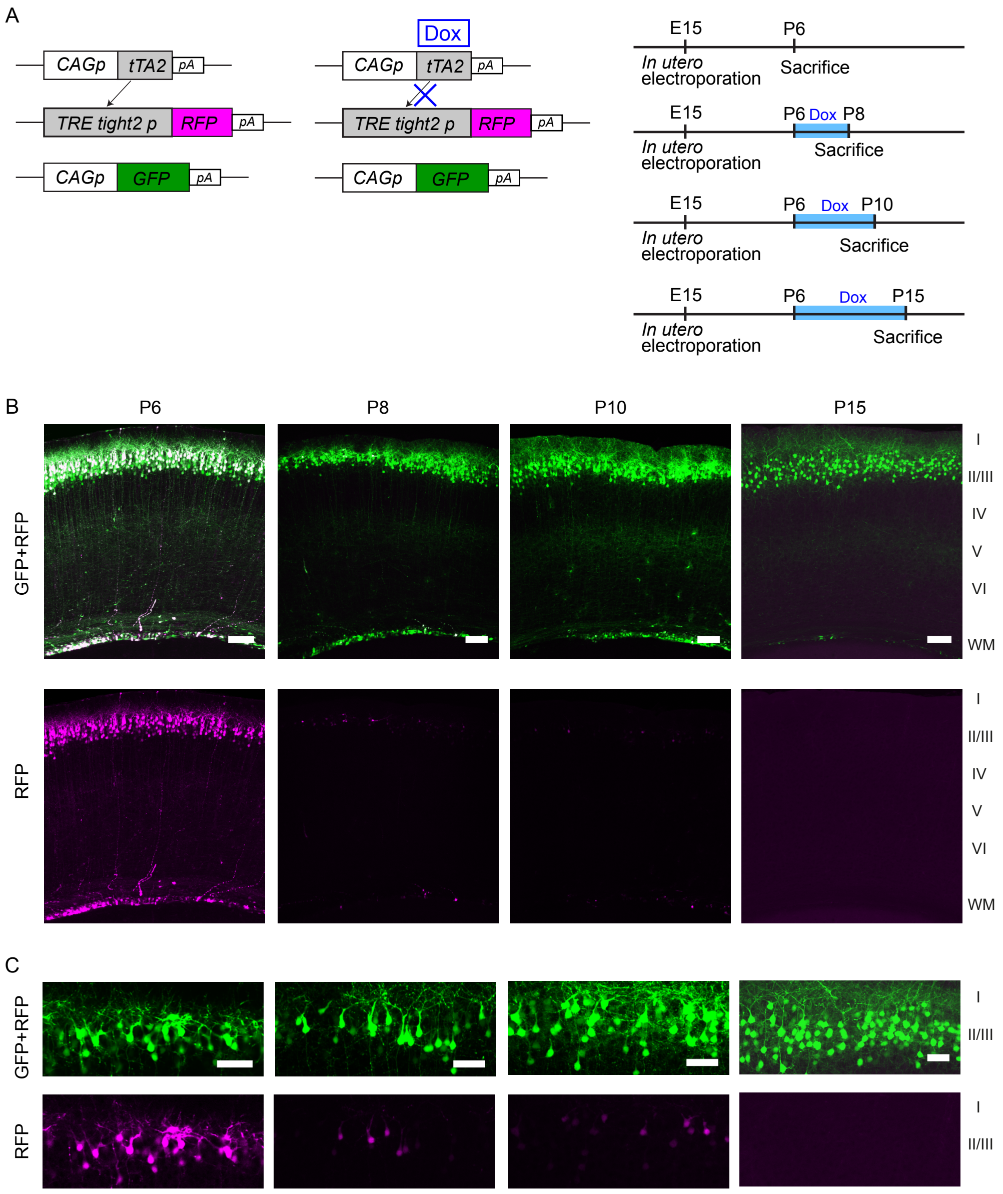


SuppleFig. 1-2

### Figure 1-figure supplement 3

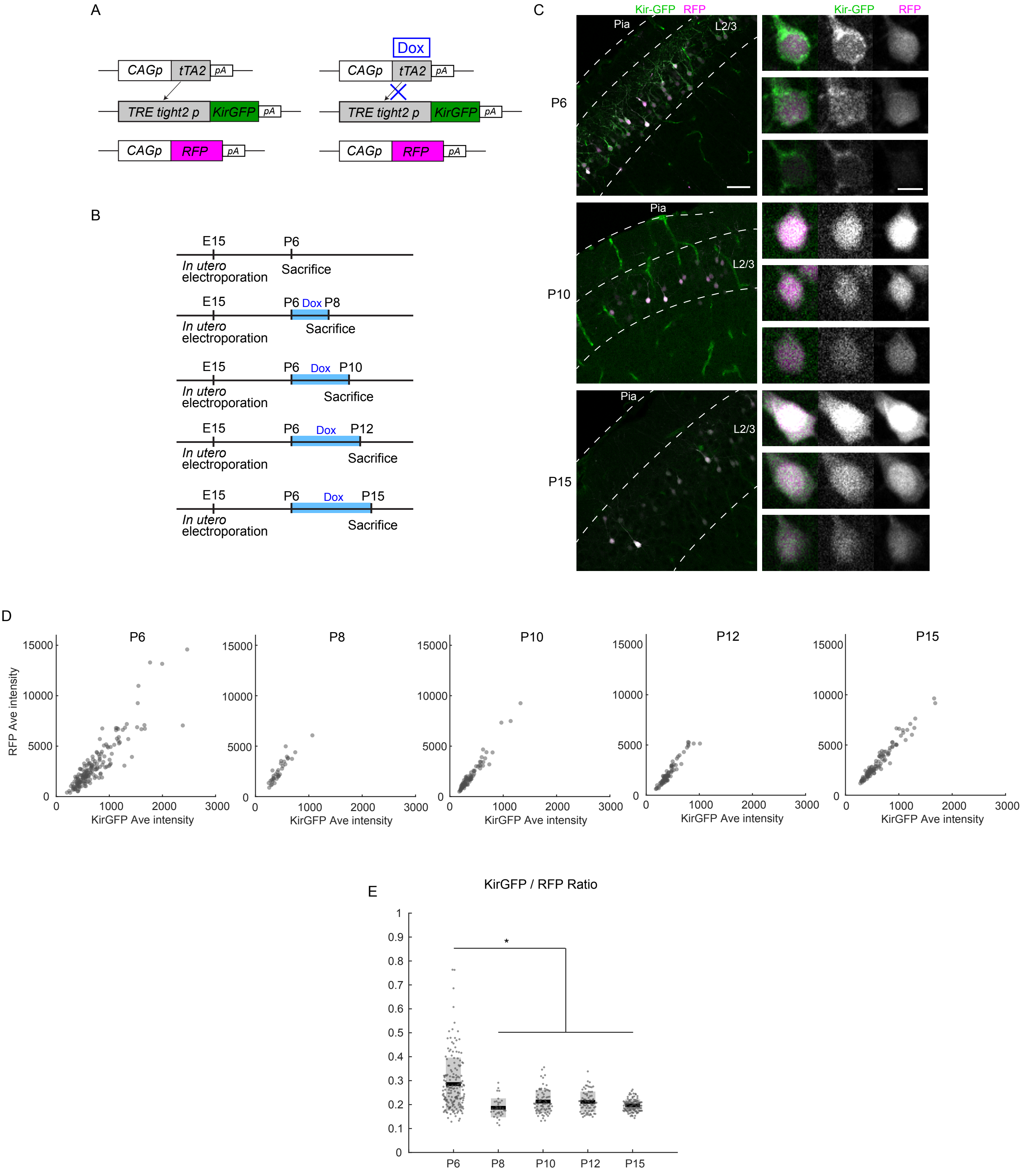


SuppleFig. 1-3

### Figure 1-figure supplement 4

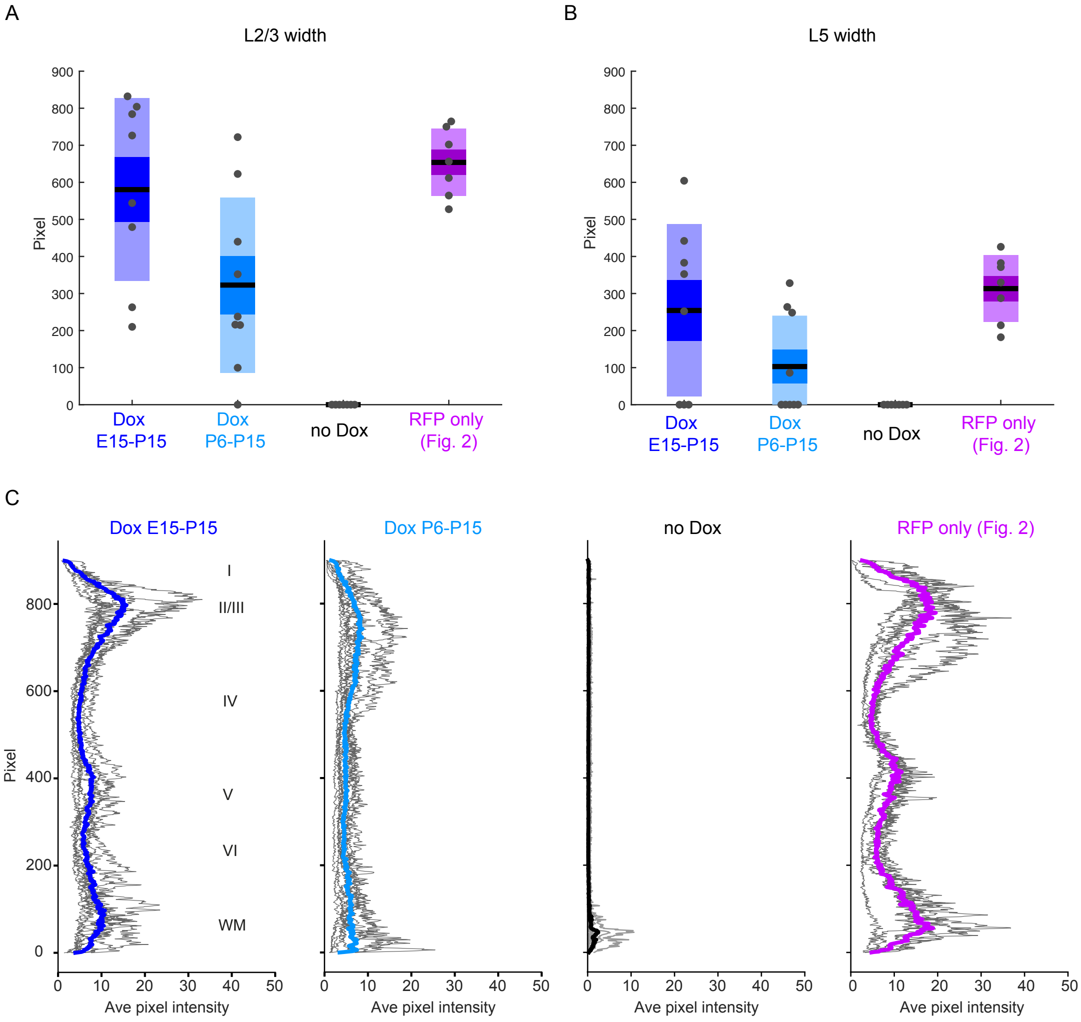


SuppleFig.1-4
